# Temporal-difference valence-partitioned Bayesian brains work out whether others are caring or uncaring

**DOI:** 10.64898/2026.05.19.725657

**Authors:** Michael Moutoussis, Adrian Frydman Laiter, Julia Griem, Donya Erfanian Delavar, Tobias Nolte, Peter Fonagy, Read Montague, Vladimir Litvak

**Affiliations:** Dept. of Imaging Neuroscience, University College London, UK; Dept. of Forensic and Neurodevelopmental Sciences, Institute of Psychiatry, Psychology and Neuroscience, King’s College London, UK; Dept. of Clinical, Educational and Health Psychology, Division of Psychology and Language Sciences, University College London, UK; Dept. of Psychology, University of Milano-Bicocca, Milan, Italy; Anna Freud Centre, London, UK; Virginia Tech University, VA, USA

## Abstract

Working out whether others care for us is crucial in personal relationships and when seeking professional help. It is often most difficult for those most in need, e.g. following interpersonal traumata. Here, we introduce a simple ‘Caring Attributions task’ and elucidate key computational mechanisms involved and, using EEG, the cortical activity representing the degree of belief that another is beneficent or maleficent. We find evidence for a new type of neurocomputational processing: valence-partitioned temporal-difference inference (TD-Bayes). This employs primary processing about latent causes, but also separate channels to represent these different valenced attributions, inspired by value-partitioned associative learning (VPAL). TD-Bayes uses slow propagation of beliefs using temporal-difference updating. These models gave a very good account of behaviour, slightly better than VPAL, but crucially, their partitioned representations have stronger, distinct representations in ERP signals. They provide a promising inroad into the understanding of how people may jump to atttibutions about caring vs. uncaring others.

## Introduction

The neurocomputational mechanisms through which people infer whether others, including health and legal professionals, may be beneficent or maleficent, is of paramount translational importance for many needs, including disputes, healthcare, post-traumatic states, paranoia and more. Helping professionals may also jump to attribute good or bad faith to help-seekers, which can be counter-productive. Psychologists talk of such categorizations as black-and-white thinking or splitting, and consider them a serious problem (Dutton, 2020). It would be very helpful to know one’s predisposition for unwarranted, polarized inferences and the brain basis of this. Towards this translational aim, we introduce a laboratory task assessing caring vs. uncaring attributions, and apply it to elucidating their computational and neural substrates.

In basic reinforcement learning (RL), costs are just negative benefits, whereas Bayesian brain theories postulate primary inference in terms of the causes of events, not of benefit and cost. We have repeatedly found that approximate-Bayesian accounts out-perform associative RL models when it comes to social cognition (Barnby et al., 2022; Hoffmann et al., 2023; Moutoussis et al., 2024). However, recent neuro-computational evidence suggests that maybe justice was not been done to associative RL, in that it is not one-dimensional (benefit-cost) but valence-partitioned (VPRL; Sands et al. (2023)). Indeed, the mesencephalon and diencephalon likely include specialized neural pathways for harm and benefit (Lawson et al., 2014; Manssuer et al., 2023). So how might higher cortical, belief-oriented processing carry out nuanced evaluation, rather than unwarranted good vs bad attributions? Does the cerebral cortex mostly compute in terms of reward and punishment, or in terms of underlying causes (e.g., people’s characters)? These computational mechanisms are as uncertain as they are translationally important.

Accounts of personal attributions based on Bayesian rationality are also challenged with respect to coherence, or integration of learning. Neural evidence indicates that the value of a particular entity A that implies B, which implies C, does not update immediately upon learning about C. Learning often has to ‘percolate’ through the states in-between, possibly by the mechanism of Temporal-Difference (TD) updating (Schultz et al., 1997; Amo et al., 2022). In TD updating, one has to perform the transition ‘A goes to B’ for the value of B to inform that of A. This contrasts with accounts of inferential, Bayesian networks which immediately pass fast messages to establish belief consistency (e.g. Moutoussis et al. (2024)).

We first asked if it is possible to reconcile the approximate-Bayesian account with temporal-difference, valence-partitioned RL. Observing that people may indeed find it natural to make discrete attributions e.g. friend, acquaintance, enemy, we propose at a theoretical solution with two components. First, we suggest that the degree of belief in each discrete attribution, or more accurately, the log of the un-normalized belief in each, can form a natural measure of value, represented within discrete neural populations. It can then be used to make decisions and express evaluations in language.

Next, we suggest that step-by-step experience is needed for these Bayesian attribution quantities to be propagated, and this ‘stepping’ can be described by temporal-difference (TD) learning. Even if I inferred in C that a beneficent process is at work, it takes going from A to B to C to learn that A is a good place to be. We call this combination TD-Bayes.

At the neural level, we hypothesized that the partitioned (log un-normalized) beliefs are a key currency for cortical representations. We expected these to be electrophysiologically detectable by EEG, both in terms of distinct but overlapping belief representations, but also in terms of the weighed prediction errors updating these beliefs. Finally, we hypothesized that personal attributions may be more strongly represented in recognized ‘social brain’ areas, particularly right temporoparietal electrodes. However with respect to our translational goals, especially supporting helpers and help-seekers to make the best attributions about each other, the nature of these mechanisms is paramount, but their social specificity strictly secondary. If people have serious problems making personal attributions, and if treatments can ameliorate this, it matters very little if some other inferences, which don’t cause problems (say, about apple trees) were alo affected. These hypotheses were pre-registered in the UCL Dept. of Imaging Neuroscience (7 March 2025; https://github.com/mmoutou/CaringAttributions/tree/main/UCL_EEG/pre_reg).

To test these hypotheses, we developped a task where participants were invited to imagine that they stayed in a number of hotels, 12 days at a time. The hotel owners (partner avatars) gave advice every day about an excursion. This turned out to be Dangerous, So-so, or Great. Caring owners were less likely to expose their customers to danger. Participants first had to guess the probability that the excursion ahead would be ‘Great’. Second, after observing the outcome, they were asked about the probability that the owner was Caring, both using a visual analogue scale (VAS). We performed a pilot online task, and, in a separate sample under EEG, both this and an impersonal, control version of the task. In the latter, people were told that apples from different apple-trees would be Disgusting, So-so, or Great (See Fig. 1 and Methods for details).

**Figure 1:**
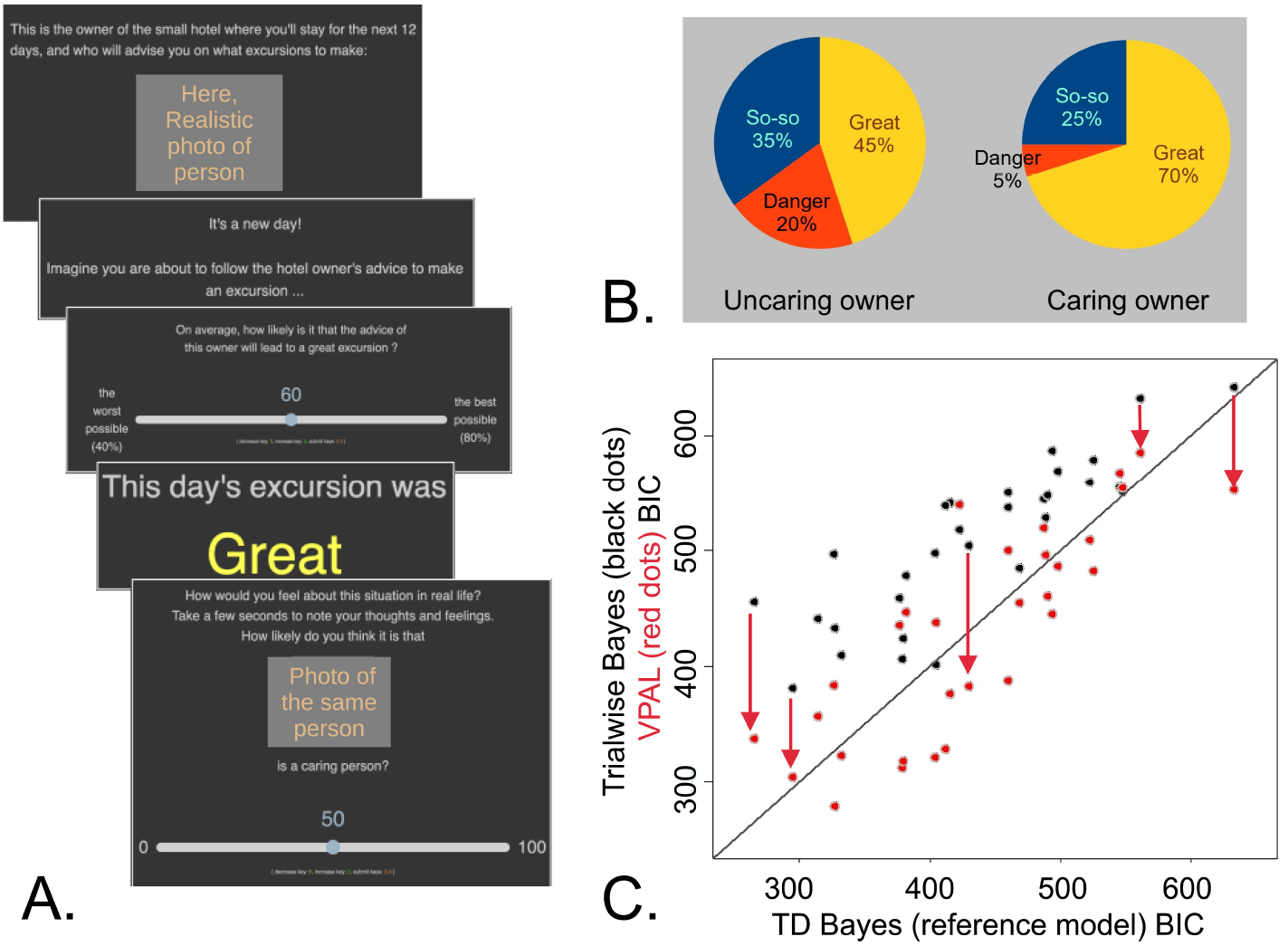
Caring Attributions task stucture and on-line results. **A**. Structure of an experimental trial. At the top, a ‘hotel owner’ is introduced with a person’s photo (here, obscured). The trial starts (state 1), then the participant is asked *how probable* it is for the hotelier’s advice to lead to a ‘great’ outing (state 2). The outcome is seen, Great, So-so or Dangerous (state 3). Then the participant is asked *how probable* it is that the hotelier is ‘Caring’ (vs. Uncaring; state 4). **B**. Participants were explicitly trained about the propabilities that beneficent vs. maleficent causes would result in great, so-so or very bad outcomes. They had to pass a comprehension test to proceed to the main task. **C**. Comparison of the TD-Bayes model with a standard, trial-wise Bayesian model (black dots), and with valence-partitioned associative learning (red dots). The standard, trial-wise Bayesian model did much worse than TD-Bayes, according to the Bayesian information criterion (BIC; Wilcox. p=3.72e-9). However, the VPAL model dramatically out-performed the standard Bayesian model, and matched the TD-Bayes model (Wilcox. p=0.44). Red arrows show examples of VPAL improving on trialwise Bayes.

## Results

### Theoretical results

The key theoretical results was that for temporal-difference updating to propagate Bayesian inferences, it suffices to encode as values the un-normalized log-beliefs corresponding to each attribution, or latent cause, in a distinct neural channel. In other words, each possible explanation for observed events can be tracked by its own neural assemblies, and a simple value-update rule can then spread how likely these explanations are forward in time, without necessitating explicit renormalization at each step. In order to retrieve the belief about the presence of one or the other latent cause, these values can be entered into a sigmoid input-output function – which is common in the brain. We call the resultant model ‘TD Bayes’, short for ‘Partitioned, Temporal-Difference propagated, Bayesian inference model’. This key result is derived in Box 1, and the model detailed in the Methods and Supplement.

#### Box 1: Partitioned Temporal-Difference propagated Bayesian inference (TD-Bayes)

Bayes rule governs updating the belief (or probability) that the professionals is caring, *P* (*C*), vs. uncaring, *P* (*U*), after an outome *o*, seen at time *t*. We rearrange Bayes’ rule to transform probabilities to value-like quantities, which we will then process via Temporal-Difference learning:

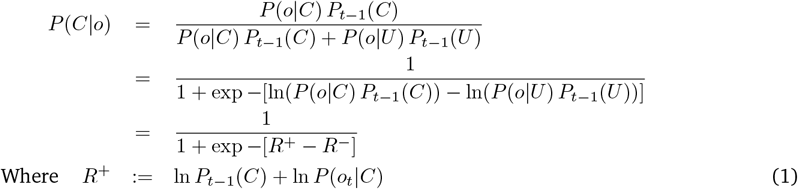

The *un-normalized log-posterior beliefs* (i.e., the log-joints) can then be *propagated as values* by standard temporal-difference learning. The change in log-belief about the professional being Caring, upon state transition *s*_*k*_ → *s*_*k*+1_, is governed by the prediction error *δ*^+^ and the learning rate *λ*^+^ :

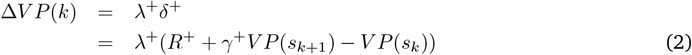

The analogous equation governs the brain’s log-beliefs about the Uncaring agent, *V N* . Estimated *V P* and *V N* can then be regressed against neural activity measures. Please see Supplement for more details.

### Pilot study

To provide pilot results and hence pre-register detailed hypotheses, on-line (Prolific Academic) modelling studies were carried out. After the task was optimized (Fig. 1 A, B), N=30 UK resident, healthy, working age (18-65) participants were tested.

We compared TD based models against a standard trial-wise Bayesian model. This simply used the likelihood function that the participants were trained on (Fig. 1 B) to sequentially update their beliefs, with forgetting decaying them to baseline (much like the *ω* parameter in Smith et al. (2022)). TD-Bayesian models had the same structure and parameters, but in addition contained valence-specific learning and discounting rates (as in Sands et al. (2023)). The crucial difference between TD-Bayes and VPAL models was that the former made use of the model (likelihood map) that the participants were taught, while the later estimated expected outcomes, not underlying causes (See Methods).

We found that TD-Bayes clearly outperformed trial-wise Bayesian models (Fig. 1 C for 29/30 partiticipants. Crucially, however, so did the VPAL model, suggesting that VPAL may be a computationally efficient way in which the brain approximates Bayesian reasoning - a hypothesis that we tested in the laboratory.

### Computational modeling

Here, we first fitted independently each of the two conditions, personal and impersonal, for each participant with each of the two best models, TD-Bayes and VPAL. We found that both models accounted for both the Expectation and the Attribution data very well, the pearson correlation between modelled and actual data being between 0.7 and 0.98. See Fig 2 A., B. for a typical example).

**Figure 2:**
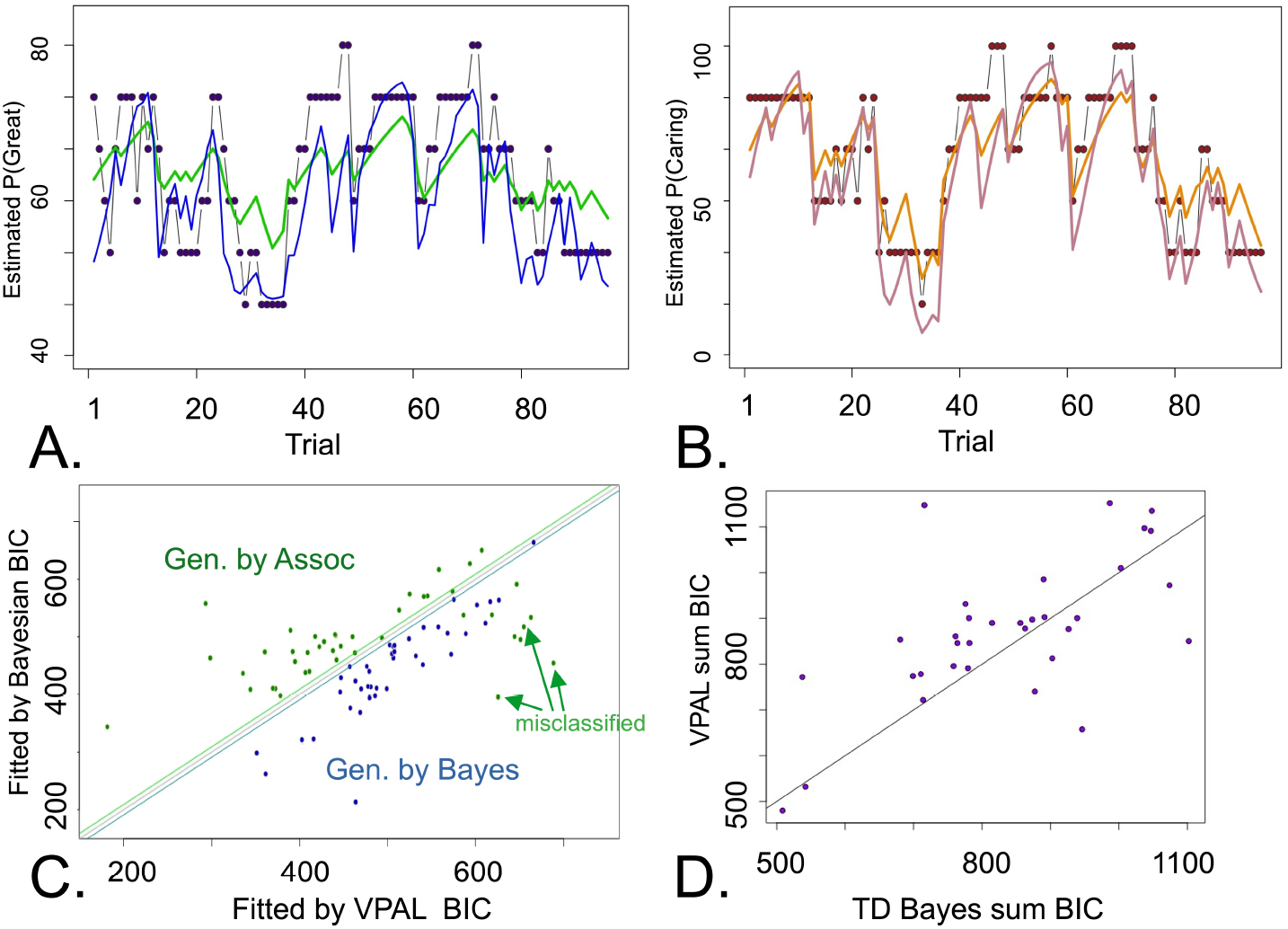
Key modelling results from the laboratory task. **A**. Typical example of the expectation (state 2 above) phase of the experiment. Black dots: experimental data, binned into 6 bins. Blue curve: TD-Bayes model expectation. Green curve: VPAL (TD Associative) model expectation, BIC=493.24. Both model preditions are highly correlated with the experimental data, *r*_*pearson*_ ≈ 0.78, *p <* 1*e−* 6. **B**. Same as A., but for the attributions phase. TD-Bayes in pink, VPAL in orange. Again both models describe the data well. Overall TD-Bayes BIC = 460.41, VPAL BIC=493.24. **C**. Model recovery study. Data produced by the fitted TD-Bayes model (blue) are never fitted better by VPAL, but 9/66 datasets produced by VPAL are better fitted by TD-Bayes (green). Green arrows indicate misclassified examples. **D**. Indicative evidence only that the laboratory participant behaviour is best accounted for by TD-Bayes, Wilcox. *p* = 0.046.)

Our hypothesis, supported by the above, being that VPAL is an efficient approximation to Bayes, we performed a model recovery analysis expecting that the two models would not be separatery recoverable. We simulated a dataset from each of the conditions of each real participant, giving 33 pts. *×*2 conditions *×*2 models = 132 datasets. Contrary to expectation, all TD-Bayes generated datasets were correctly classified (fitted better by their own class of model) and only 9 of 66 VPAL generated datasets were misclassified.

We then compared model fits, and found that experimental data across conditions were more often best-fitted by the TD-Bayes rather than VPRL model – but, unlike for simulated datasets, the evidence was modest. 24/33 cases were best fittd by TD-Bayes, 9/33 cases by VPAL, Wilcox. *p* = 0.046, Fig. 2 D. This suggested that neural data would be useful to disambiguate participants’ cognitive mechanisms.

Finally, we tested whether participants preferentially used TD-Bayes for personal rather than impersonal attributions. We found no evidence for this (wilcoxon *p* for the advantage of TD-Bayes over VPAL in personal context – same for impersonal context = 0.525).

### Condition-based EEG analysis

We first examined averaged Event Related Potentials (ERPs) for each of two factors, sociality (personal vs. impersonal) and outcome (bad - mid - good), i.e. 2*×* 3 levels. Several ERP features were discerned, corresponding closely to published results for stages of cognitive processing in complex learning tasks. ERP activity was mostly centered near electrodes Pz (Fig. 3 A) and Fc4 (Fig. 3 B).

**Figure 3:**
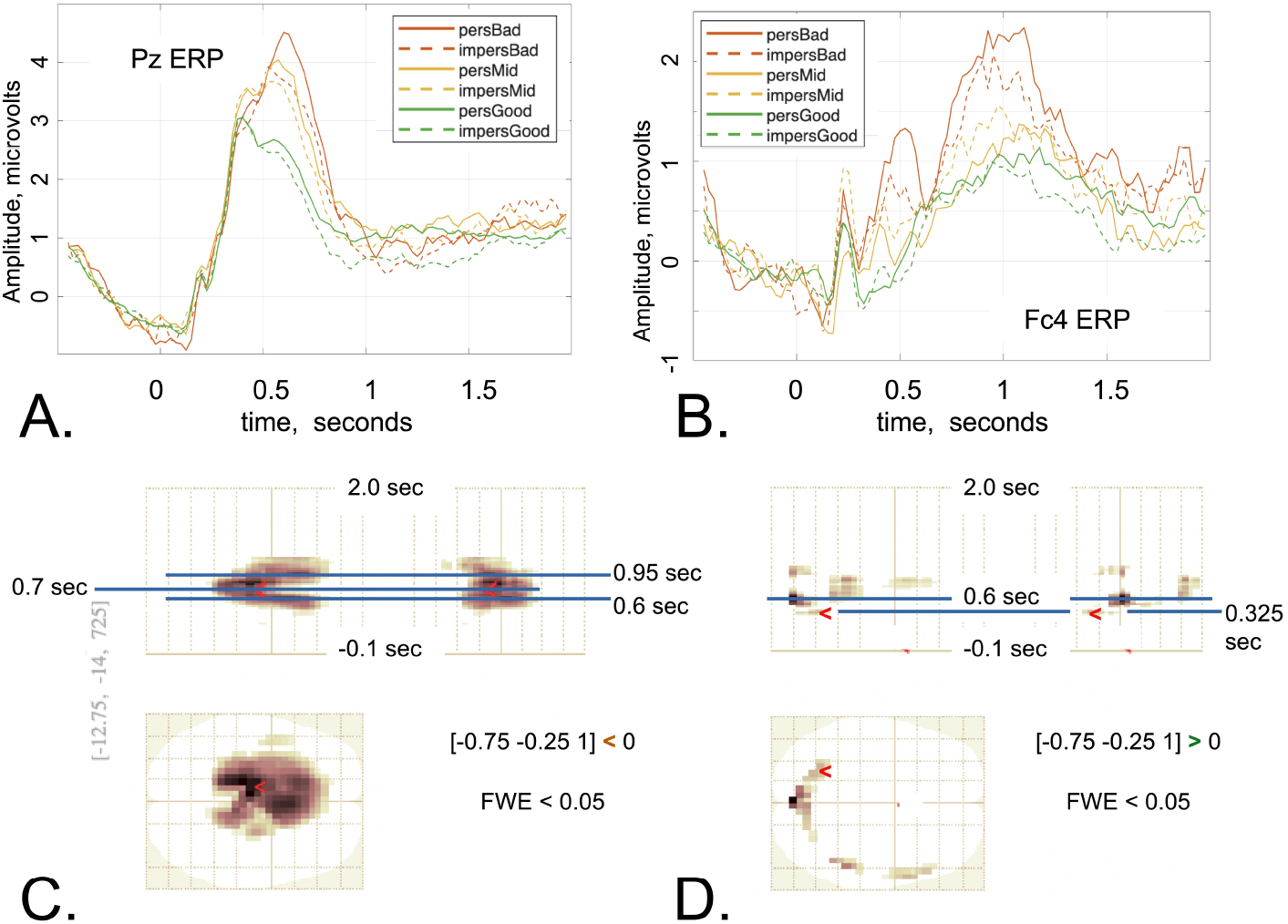
ERP analysis by experimental condition **A**. Parietal (Pz) ERP, showing negatively slopping ‘preparatory’ baseline, main peak at about 0.6 sec and late positive plateau. **B**. Right frontal (Fc4) ERP, additionally showing early evaluative peak at around 0.25 sec, and a large late-processing peak at 1 sec, as well as features analogous to Pz **C**. Good - NotGood outcome voltage difference *<* 0 contrast, showing complex perivertical activity 0.5 →0.7 →0.95 sec. **D**. Good-NotGood outcome voltage difference *>* 0 contrast, with temporal, temporo-parietal and iniac clusters. Marker at the earliest positive cluster, with *p*_*FW E*_ = 0.001

We then performed formal statistical analysis using SPM25, as per pre-registered analyses (UCL Dept. of Imaging Neuroscience, 2025; github.com/mmoutou/CaringAttributions/tree/main/UCL_EEG/pre_reg), examining the main effects of outcome, sociality and their interaction. As participants were instructed that ‘so-so’ outcomes were somewhat more common if their partner was Uncaring, we weighted outcomes *a priori* as *c* = [*−* 0.75 *−* 0.25 + 1], bad-mid-good. Several scalp voltage*×* time significant culsters were present for the outcome contrasts (FW E *<* 0.05, Fig. 3), especially NotGood *>* Good (Fig. 3 C).

Significant clusters were also present for the Good *>* NotGood contrast, albeit with a different topography. The timing of all contrast clusters roughly corresponded to the key peaks of the ERP (blue lines, Fig. 3 C,D; cf. A). This included an early left posterior cluster with peak at 0.325 sec.

However, there were no significant cluters for the main effect of sociality, nor for a sociality-by-outcome interaction.

### Modeling-based EEG analysis

We derived participant-specific parametric regressors from the TD-Bayes and VPAL models, for both weighed prediction errors (VPAL *λδ*) and the corresponding log-belief updates, Δ*V* and positive/Caring and negative/uncaring values (*V P, V N*, Box 1). We applied these to the outcome delivery epoch in the task, using them to test the preregistered hypotheses that a) distribution of prediction errors would differ per valence, aversive to appetitive, and sociality, impersonal to personal and b) the neural representation of appetitive and aversive values would differ.

We found no significant clusters for any of these regressors for the VPAL model, but we found significant clusters both for belief update and for partitioned values for the TD-Bayesian model. With respect to belief update, we first found an extended significant cluster (*p*_*FW E*_ *≈* 0.000) containing with three peaks (*xy* = (*−* 4, *−* 19), *t* = 0.65*s*; *xy* = (*−*17, *−*41), *t* = 0.675*s*; *xy* = (*−* 8, *−*9), *t* = 0.775*s*, all *p*_*FW E*_ *≈*0.000). This pertained to the Caring belief update, and negative voltage (Fig. 4A). For the same belief, but positive voltage, we found two significant but smaller clusters with similar timing but right temporal distribution (*p*_*FW E*_*≈* 0.000 and *p*_*FW E*_0.003). They contained several peak voxels significant at *p*_*FW E*_ *<* 0.05, the main two ones being at *t* = 0.525*s, p*_*FW E*_ = 0.011 and *t* = 0.675*s, p*_*FW E*_ = 0.008 (Fig. 4B). To better visualise this neural activity, we plotted the belief-update regressor contrast image, simply averaged over participants, at *t* = 0.525*s*, revealing the positive and negative clusters (yellow and blue, Fig. 4C). Finally, we plotted the time-course of the this average-over-participants near the negative-peak voxel, indicating that the FW E-corrected peaks are likely underpinned by more extended neural representation of these computational regressors (Fig. 4D). We then turned to the Uncaring belief update, and here we found a significant positive-voltage, left temporal cluster. Here, neural representation appears to ‘ramp up’ during the epoch examined (Fig. 5A, B).

**Figure 4:**
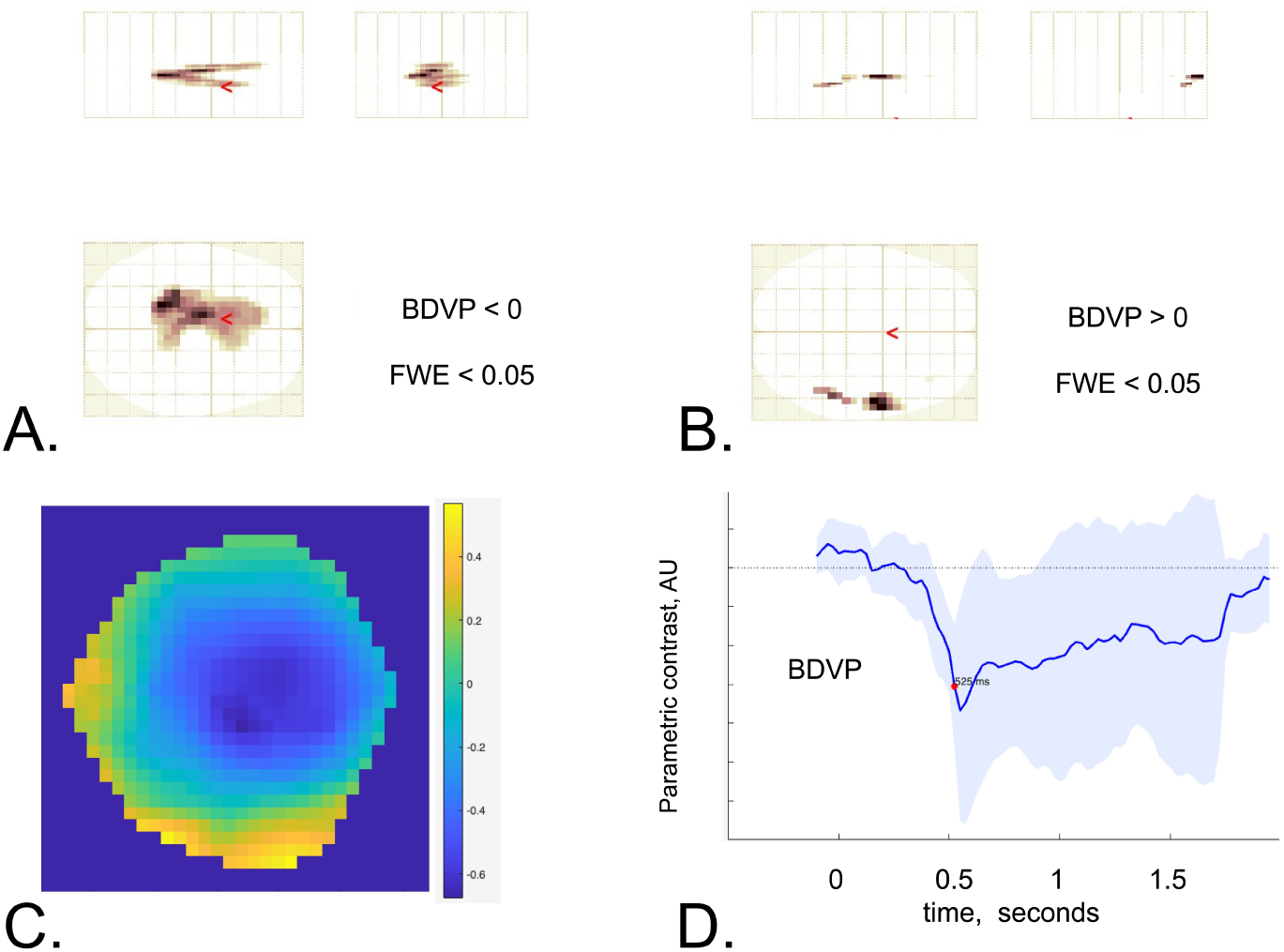
Parametric regressors for update of beneficence belief **A**. Regression of the Beneficent log-probability update, Δ*V P*, contrast *<* 0. Activations resembling the topograpy of the Pz ERP are seen, including a frontal midline cluster (from 0.6 sec) and a parietal one. **B**. The opposite contrast for the same regressor shows right temporal significant cluster **C**. Average of the beta (Δ*V P* regressor) contrasts in A. over participants at the time of the first significant negative-peak voxel, 525 mSec, showing both the midline negativity and temporal positivity **D**. Time-series of a 3x3 cluster average of the same regressor around the negative-peak voxel of C. Red dot is time of C.

**Figure 5:**
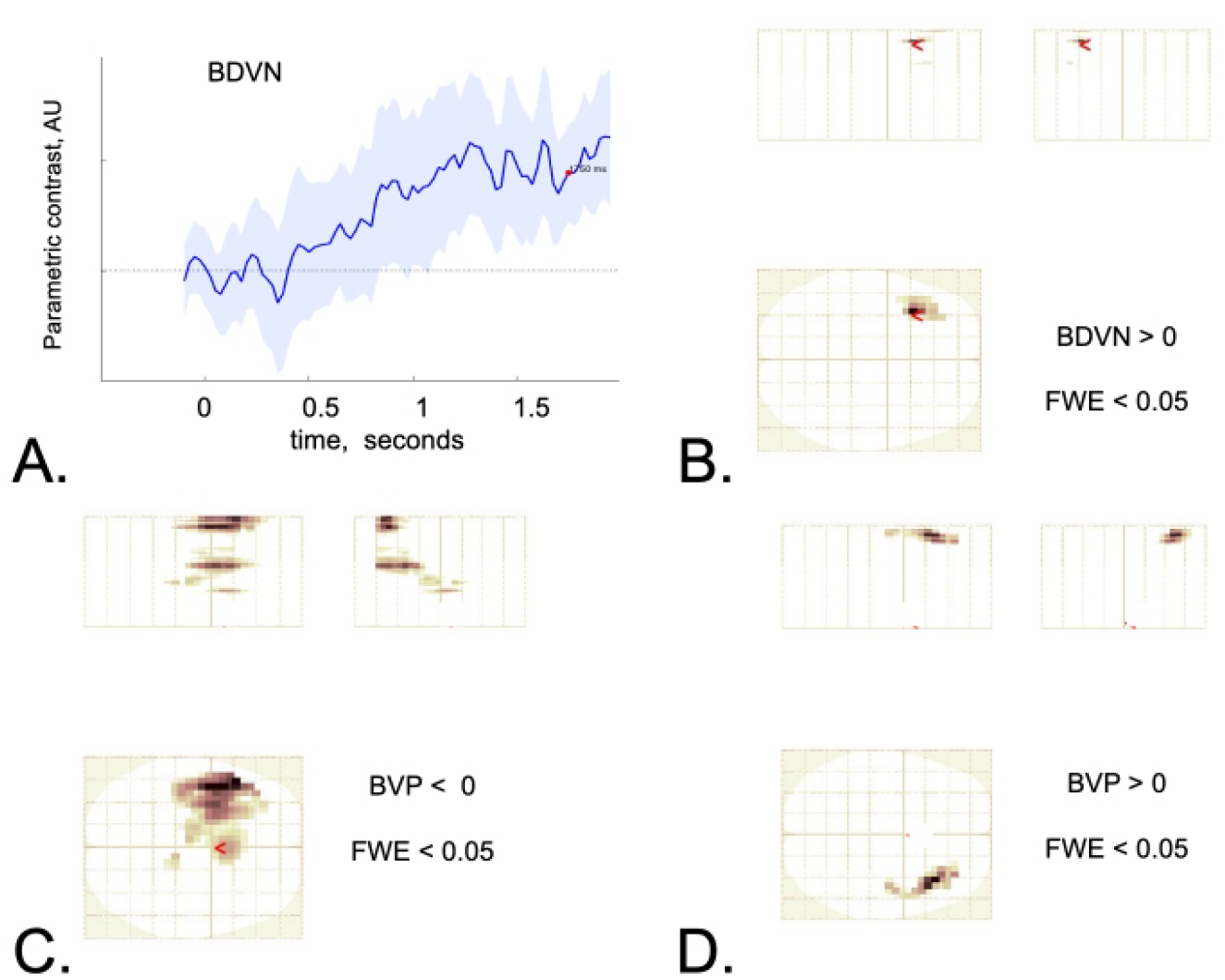
Update of maleficence beliefs, and representation of beneficence value. **A**. Time-course of averaged 3x3 voxels at a left temporal location that emerges as a significant cluster, for the update of the maleficence belief. Red dot marks the time of the peak of the significant cluster in question. **B**. The left temporal cluster, whose peak shows the time-course in A. This is the domain of positive voltage. **C**. Complex representation of the beneficence value (log-belief, *V P*) in negative voltage. There are three main peaks, one near Fz at around 0.5*s* and two more left-lateral ones at 1*to*2*s* **D**. The same regressor, *V P* has a positive-voltage correlate at a right temporal late in the epoch.

We also found significant clusters associated with the Caring belief regressor, but no significant clusters for the Uncaring belief. The Caring regressor was represented by negative voltage within three clusters, one around 0.525 s near Fz (*p*_*FW E*_ *≈*0.000, *xy* = (4, 13)) and two much more extended left temporal ones, spanning from 1 *s* to 2 *s* approx. (Figure 5C; *p*_*FW E*_ *≈*0.000 to 0.025, *xy* from (*−*42, 13) to (*−*47, 18)). We also found a positive voltage right-temporal cluster late in the epoch (Fig. 5D, *p*_*FW E*_*≈* 0.000, peak *xy* = (38, 24)).

None of the significant results above showed modulation by sociality – that is, there were no significant clusters for the interaction of the social/non-social condition with belief-update or belief-representation regressors.

## Discussion

In this work, we asked how healthy people make attributions about whether others are caring or not. Attributions about whether others are caring or uncaring are central to help-seeking and engagement, and thus highly relevant to the helping professions. Our computational and EEG results inform a mechanistic account of these attributions, in neurocomputational and electrophysiological terms. In pilot work using a novel task, we found that associative *valence-partitioned attribution learning* (VPAL) out-performed previously successful Bayesian models, challenging these previous results. We asked whether it is possible to reconcile neurobiological evidence for temporal-difference based updates of neural representations with an inferential account of attributions. Hence, we theorized that un-normalized posterior inferences about key features of the enviroment are propagated within neural channels that each use temporal-difference updating. We call this the TD-Bayes model. TD-Bayes gave a slighty better account of our behavioural data than VPAL, but its underlying belief-related measures correlated much more clearly with cortical EEG activity than those of VPAL. We found no significant differences between personal and impersonal versions of our task, as to either their computational or neural underpinnings.

The success of temporal-difference updating to account for behaviour in our task, which probes behaviourally two of four within-trial states, is remarkable. It suggests that, even when people can perform approximate Bayesian inference, they update their beliefs incrementally from prediction errors over time, rather than establishing global consistency at once. In addition, the relative success of the VPAL model in describing behaviour while failing to capture the neural data means that current valence-partitioned associative models may be too flexible. Hence, they can ‘curve fit’ behaviour without capturing its underpinning cortical dynamics, much like the epicyclical model of the planets predicted their positions without capturing planetary dynamics (Freeth et al., 2021).

In the simple inferential – rather than associative – partitioned model that we put forward, TD-Bayes, it is beliefs about beneficent and maleficent (social) causes of events that are processed in parallel. This leads to the prediction that both the topography and the timing of representations of beneficent and maleficent causes, including social causes, must differ across the cortex. EEG evidence supported this prediction.

The neural data spoke to valence-partitioned attributions. Sociality*×* Outcome ERP analyses showed widespread activity significatly associate with outcome goodness, from 0.325 sec for good - bad outcomes, to *∼* 1 sec later. The pattern is consistent with previous accounts of appetitive pathways signaling faster (Sands et al., 2023), possibly as they involve fewer synapses. Parametric analysis showed that the main, P3b cluster represented the update of beliefs about beneficence, according to the TD-Bayes model, and not the VPAL one. Update of maleficence beliefs was also detected, but at a much later, left frontotemporal cluster, outside the main P3b complex. We also found strong, complex, temporally extended representation of the beneficence beliefs themselves (rather than their updates), extending to the boundary of the epoch. Speculatively, this may be because participants knew that the up-coming rating would be ‘What is the probability that the agent is Caring?’ (or ‘soil is good?’). In line with the ERP results, and against our pre-registered hypotheses, we found no evidence that the social condition may involve different spatio-temporal representations of beliefs, nor their updating.

We argue that in scenarios resembling the one we tested, the social importance of cognition is high, but its social specificity is of very little interest to help-seekers and professionals, who engage with beliefs that others are harmful. They are interested in the substrates of social cognition, not whether the same substrates may also be used for purposes which do not trouble them. The low relevance of social specificity here stands despite vast evidence, including our own modeling (Garvert et al., 2015; Barnby et al., 2022), that other aspects of cognition are socially specific. From an evolutionary point of view, it would be highly disadvantageous if social cognition didn’t recruit generically optimized structures computing beneficence and maleficence. However, the question of social specificity is of interest to the neuroscience community. Hence, we compared the cognitive and neural substrates of our socially framed task with a non-social control. Against the specificity hypothesis, and consistent with the generically optimized resource one, we found no computational or neural specificity of the valence-partitioned mechanisms. Still, this does not mean that no such socially specialized systems exist. In addition, our within-participant design rendered it likely that cognitive cross-contamination took place. In a microcosm of the evolutionary argument just presented, the mathematically similar nature of the social and control tasks would encourage participants to use whatever inferrential solution they discovered, across conditions.

### Limitations and further research

Our study has several limitations. First, using categorical caring / uncaring inference might itself push participants to use immature, good-bad, rather than integrated thinking (Kernberg, 1975). However, in previous work we found substantial evidence that people do use underlying polar categories when these are not provided by the researchers, only then integrating them in a ‘shaded’ manner (Lau et al., 2024). Second, in order to examine ordinary, peacetime inference our scenarios included few (though very salient) very negative outcome trials, and this is likely to have impaired our ability to detail the neural representation of maleficence and its updates. Third, despite constructing likelihood maps for face validity (Fig. 1), training participants on these maps sacrifices ecological validity and individual variation for experimental control. Future studies should be adequately powered to detect individual models of caring and uncaring others, especially in high-maleficence contexts, and how people resolve uncertainty about these. Future studies should also address whether the mechanisms we describe, or alternative ones, are engaged in interactive helper – help-seeker interactions. Although not relevant to our translational motivation, researchers interested in the fine grain of social specificity of brain inference should study participants’ own social vs. non-social models (rather than train them on one, like we did.). They should also use between-participant designs to reduce cross-condition cognitive contamination.

## Conclusion

In order to elucidate basic neurocomputational mechanisms behind how people infer whether others are caring or uncaring, mechanisms very important for helper – help-seeker relationships, we introduced a novel ‘Caring attributions task’. We found substantial behavioural and EEG evidence that people infer about others on the basis of a model of their behaviour, that temporal-difference updating captures how they propagate their inferences through extended cognitive models, that they encode their beliefs in valence-partitioned channels, and that these carry distinct EEG signatures. We found no evidence for the social specificity of the mechanisms involved. These findings lay key foundations for the future study of constructive vs. counter-productive attributions by receivers (and providers) of help within caring relationships.

## Methods

### Participants and Materials

We recruited two groups of participants. The first was a pilot on-line group of N=30 UK residents with no current history of neurological or psychiatric disorders, learning impairments or substance use disorder, between the ages 18 to 65. The main, EEG study group had N=33, who also fulfilled the same criteria. 21 reported their sex at birth to be female, and 12 male. They were diverse, from six ethnicity groups, the most common being ‘non-UK white’ (n=10). The most common occupations were ‘student’, n=18, and professional (engineer, accountant etc; n=7). Participants were paid 14 pounds sterling per hour with no performance bonus. All gave written informed consent, UCL Ethics number: 19601/004.

The task was coded and delivered by the first author using PsychoPy (Peirce et al., 2022) and was near-identical for the online and EEG variants. EEG was recorded using a BrainAmp MR plus system with actiCAP slim electrodes (BrainAmp MR with ActiCap, Brain Products, Gilching, Germany). Data were acquired using BrainVision Recorder software. Computational modeling of behaviour was coded in R (Giorgi et al., 2022) and matlab (matlab, 2019). All EEG analysis was carried out in SPM25 (Tierney et al., 2025).

### Computational modeling

The simplest, trial-wise Bayesian model includes several concepts that the more complext models inherit (Table 1; See Supplement for equations). The first is a parameter representing the strength of baseline positive expectation that participants have. In all Bayesian models, this is the baseline probability that others will be Caring. All models also have a block learning rate, for updating baseline positive expectation between blocks of trials, i.e. from one on-screen partner to the next. Next, all models have a decision noise parameter, that sharpens or blunts the probability of implementing one’s preferred self-report decision. All models also include two parameters modulating self-reports that one will experience a ‘great’ outcome, and that one’s partner is Caring. These do not affect underlying cognition, but are mainly self-report biases, e.g. social desirability biases. A lapse-rate, motivation-independent noise parameter is also always included. Finally, the all Bayesian models include a forgetting or volatility parameter. The trial-wise Bayesian model then starts from baseline positivity beliefs; uses the likelihood map (Fig. 1 B) to estimate the probability of Great outcome and reports this, influenced by lapse rate and decision noise. Then, if not already at baseline, drifts its beliefs back towards the baseline, influenced by the forgetting rate. It observes the outcome, and updates its beliefs according to Bayes rule; and finally, it reports its beliefs about the partner being Caring, again subject to noise. The cycle then repeats.

**Table 1:**
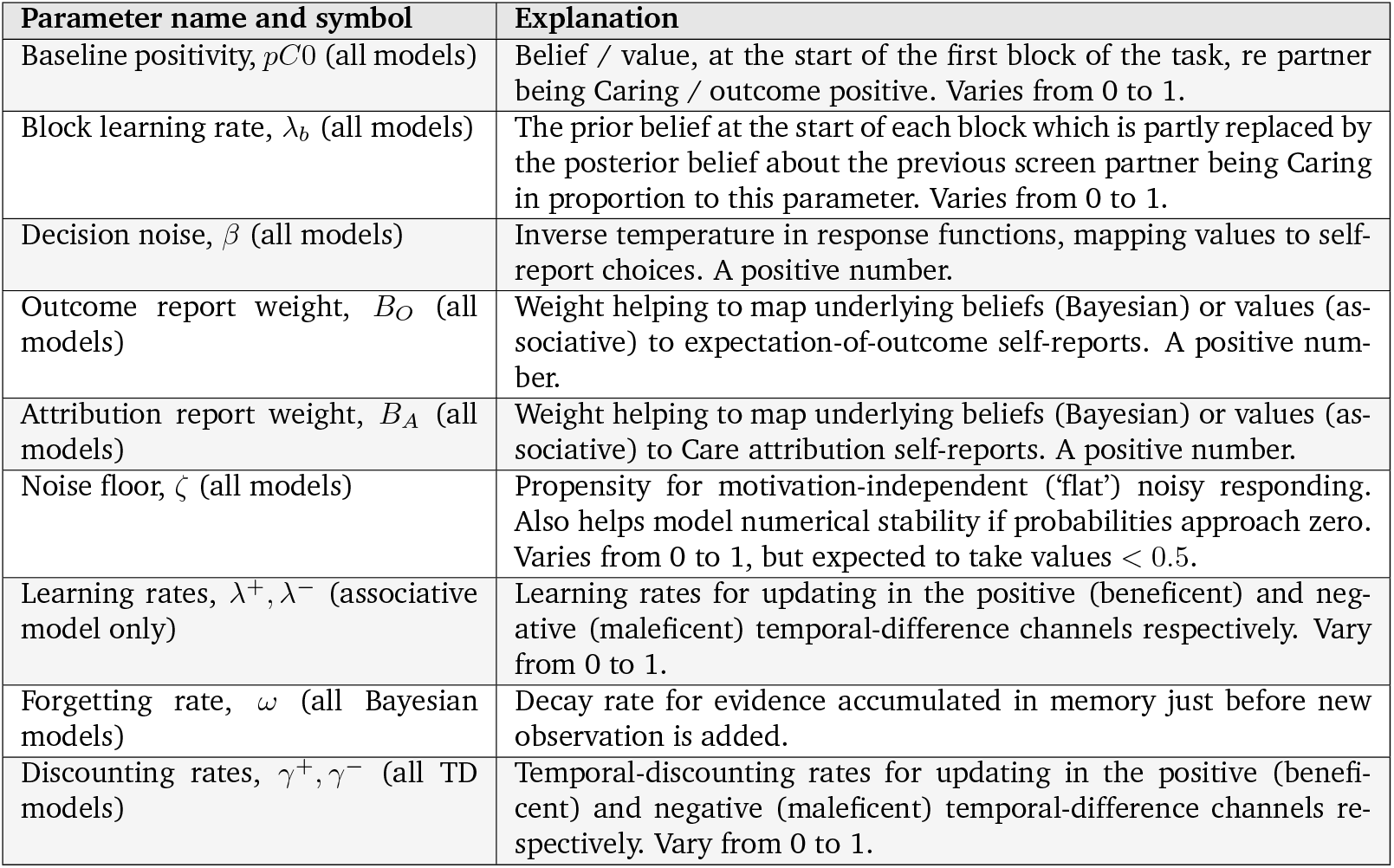
Parameter table.

| Parameter name and symbol | Explanation |
| --- | --- |
| Baseline positivity, $pC0$ (all models) | Belief / value, at the start of the first block of the task, re partner being Caring / outcome positive. Varies from 0 to 1. |
| Block learning rate, $\lambda_b$ (all models) | The prior belief at the start of each block which is partly replaced by the posterior belief about the previous screen partner being Caring in proportion to this parameter. Varies from 0 to 1. |
| Decision noise, $\beta$ (all models) | Inverse temperature in response functions, mapping values to self-report choices. A positive number. |
| Outcome report weight, $B_O$ (all models) | Weight helping to map underlying beliefs (Bayesian) or values (associative) to expectation-of-outcome self-reports. A positive number. |
| Attribution report weight, $B_A$ (all models) | Weight helping to map underlying beliefs (Bayesian) or values (associative) to Care attribution self-reports. A positive number. |
| Noise floor, $\zeta$ (all models) | Propensity for motivation-independent ('flat') noisy responding. Also helps model numerical stability if probabilities approach zero. Varies from 0 to 1, but expected to take values $< 0.5$. |
| Learning rates, $\lambda^+, \lambda^-$ (associative model only) | Learning rates for updating in the positive (beneficent) and negative (maleficent) temporal-difference channels respectively. Vary from 0 to 1. |
| Forgetting rate, $\omega$ (all Bayesian models) | Decay rate for evidence accumulated in memory just before new observation is added. |
| Discounting rates, $\gamma^+, \gamma^-$ (all TD models) | Temporal-discounting rates for updating in the positive (beneficent) and negative (maleficent) temporal-difference channels respectively. Vary from 0 to 1. |

The Associative, VPAL model shares all the constructs of the simplest Bayesian model (baseline positivity and its updating between blocks, noisy, biased responding, etc) – except a separate forgetting parameter and, of course, making use of the likelihood map. However, it introduces Temporal Difference (TD) updating, closely following Sands et al. (2023). That is, what is learnt upon making an observation does not form knowledge automatically available to all the stages of the next trial. Instead, knowledge is propagated only step-by-step, as one moves from an un-updated state to the next. This necessitates separate TD learning rates and discount parameters (cf. Box 1). Crucially, it uses partitioned values, so that positive and negative outcomes are accounted for and propagated separately.

Finally, the TD-Bayesian model shares the general features of both the above. Upon observing the stimulus, the TD-Bayes agent uses the likelihood map to update its beliefs about others being Caring vs Uncaring. Then, it uses temporal-difference updating to propagate this knowledge. However, unlike the VPAL model, what’s propagated here isn’t information about gains and losses, but beliefs about others being caring or uncaring (see Box 1). The TD-Bayes model has one fewer parameter than VPAL, as it retains the forgetting of trial-wise Bayes, but does not need associative learning rates to process observations.

### Model-fitting

We used maximum-a-posteriori fitting with uninformative priors, to maximize sensitivity to detect correlations between modelling and non-modelling measures (Moutoussis et al., 2018). After a first-pass fit to the sample, we used an initial condition grid with each parameter initiated from the first and third quartile of the sample distribution. This was effective in minimizing local-optimum errors. We used the Bayesian Information Criterion (BIC) at the individual level to account for model complexity.

### EEG Analysis

ERP analysis was based on Zhang et al. (2023). Each participant’s EEG was filtered between 0.2 and 10 Hz. Data was then inspected visually and obvious bad channels marked by hand. First-pass eyeblink detection was carried out using channel Fp1. One-second epochs around these eyeblinks were constructed, and blink ERPs were constructed by robust averaging. Then, the spatial blink pattern was extracted using singular value decomposition. The data was down-sampled to 40 Hz and epoched from -450 mSec to 2000 mSec around outcome onset. Using the spatial blink pattern, we removed blink related activity by Signal Space Projection in SPM25. For purposes of display and quality control, data was then high-pass filtered at 0.2 Hz again, and robust averages were obtained for sociality*×* outcome conditions. In contrast, for purposes of statistical analysis, the pre-processed data was converted to images.

Images were first analysed using a ‘flexible factorial’ design with two factors, outcome with 3 levels and sociality with 2 levels. Each of these six were entered separately into the design matrix. As described in **Results**, we had specific hypotheses regarding valence, hence we used the contrast [*−*0.75, *−* 0.25, 1] and its negative over bad, middle and good outcomes. The main effect of sociality was the t-contrast [*−*1, *−*1, *−*1, 1, 1, 1, ]*/*3 for *personal*-*impersonal* (also its negative), the main effect of outcome [*−*0.75, *−* 0.25, 1, *−*0.75, *−*0.25, 1], their interaction was [0.75, 0.25, *−*1, *−*0.75, *−* 0.25, 1], etc. At second level, no explicit masking was used. Significance for clusters detected was taken to be family-wise-error (FW E) level 0.05.

For the parametric analysis, first, a pair of regressors was formed from the positive-valence and negative valence values (or beliefs) from each model at the time of outcome. We also formed an analogous pair for the weighed-prediction error at the time of outcome. For associative models, the weights in question were constant, but for Bayesian models, the weight was a balance of prior precision and likelihood precision.

## Acknowledgements

We would like to thank Terry Lohrentz for his support with this project. The Department of Imaging Neuroscience is supported by a platform grant from the Welcome Foundation. Read Montague is supported by the Wellcome Trust and the Red Gates Foundation. Peter Fonagy is supported by a National Institute for Health Research (NIHR) Senior Investigator Award (NF-SI-0514-10157).

## CReditT

1. Michael Moutoussis: Conceptualization, methodology, software, formal analysis, investigation, resources, data curation, writing all drafts, supervision of AFL and DED, project administration.
2. Adrian Frydman Laiter: Conceptualization, methodology, formal analysis, investigation, resources, data curation, editing.
3. Julia Griem: Investigation, editing.
4. Donya Erfanian Delavar: Investigation, formal analysis, data curation, editing.
5. Tobias Nolte: Investigation, editing.
6. Peter Fonagy: Investigation, editing, supervision
7. Read Montague: Investigation, editing, supervision
8. Vladimir Litvak: Conceptualization, methodology, softwared, resources, editing, supervision of all aspects of EEG.

## Supplement

### Task details and the impersonal task

The impersonal and personal tasks were mathematically identical, and were as close as possible in terms of presentation. However, it was stressed in the instructions that in the personal task, the hotelier being caring or uncaring determined the likelihood function; whereas in the impersonal version (the ‘apple trees task’), outcomes depended on the quality of the soil. Thus, the impersonal task did not have the inter-personal salience of the social task. Fig. S1 shows an example screen-shot from the ‘apple trees’ version.

In the impresonal version, the cover story had as follows: Participants were told that they were apple enthusiasts, who cared about apple trees. They visited a number of hotels, like in the personal task - but this time they were concerned with the hotel’s orchard. Their objective was to work out whether the soil was conducive to great apples, or was too acidic. They experienced the same number of trials, or apples per orchard (12), as in the personal task. The apple could taste Disgusting, So-so or Great. This was meant to approximate as closely as possible the social task outcomes, but make sense for apple tasting. They were trained on exactly the same likelihood map, using the same pie charts as Fig. 1B. They were told that the orchard owner was always as good a farmer, on average - this had nothing to do with caring, so they should focus entirely on whether the soil was good or too acidic to explain their observations. We simply told participants that we wanted to compare how they thought about people vs. apples, and compare the brain signals involved.

### Model formulation

Here we describe the models and their equations step-by-step. Please see Table 1 in the main text for parameter descriptions.

#### Trialwise Bayesian model

The assumed belief that the ‘hotel owner’ is caring before seeing the outcome of first trial is given by the free parameter *pC*0:

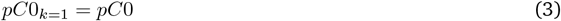

In general, between blocks (i.e. hotel owners) *k −* 1 and *k*, this baseline belief is simply updated by an admixture of what was learnt about the last hotel owner’s character in the task. We denote the belief in Caring at the end of the previous block as *pC*_*T*_, and write:

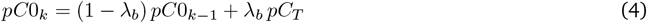

This is the simplest possible rule, which does not consider precision-informed belief updates.

Within each block volatility (equivalently, forgetting) drifts beliefs back to the prior *for that partner*. For hotel owner (i.e., trial block) *k*, trial *t*, we notate *pC*_*t−*1_ the belief at the end of excursion (i.e., trial) *t −* 1 and write:

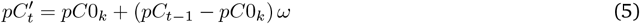

When asked ‘what is the probability that the outcome of this excursion will be Great?’, Bayesian participants marginalise over their beliefs in caring partner *C* or uncaring, *U*, according to the known likelihood function (Fig. 1B):

**Figure S1:**
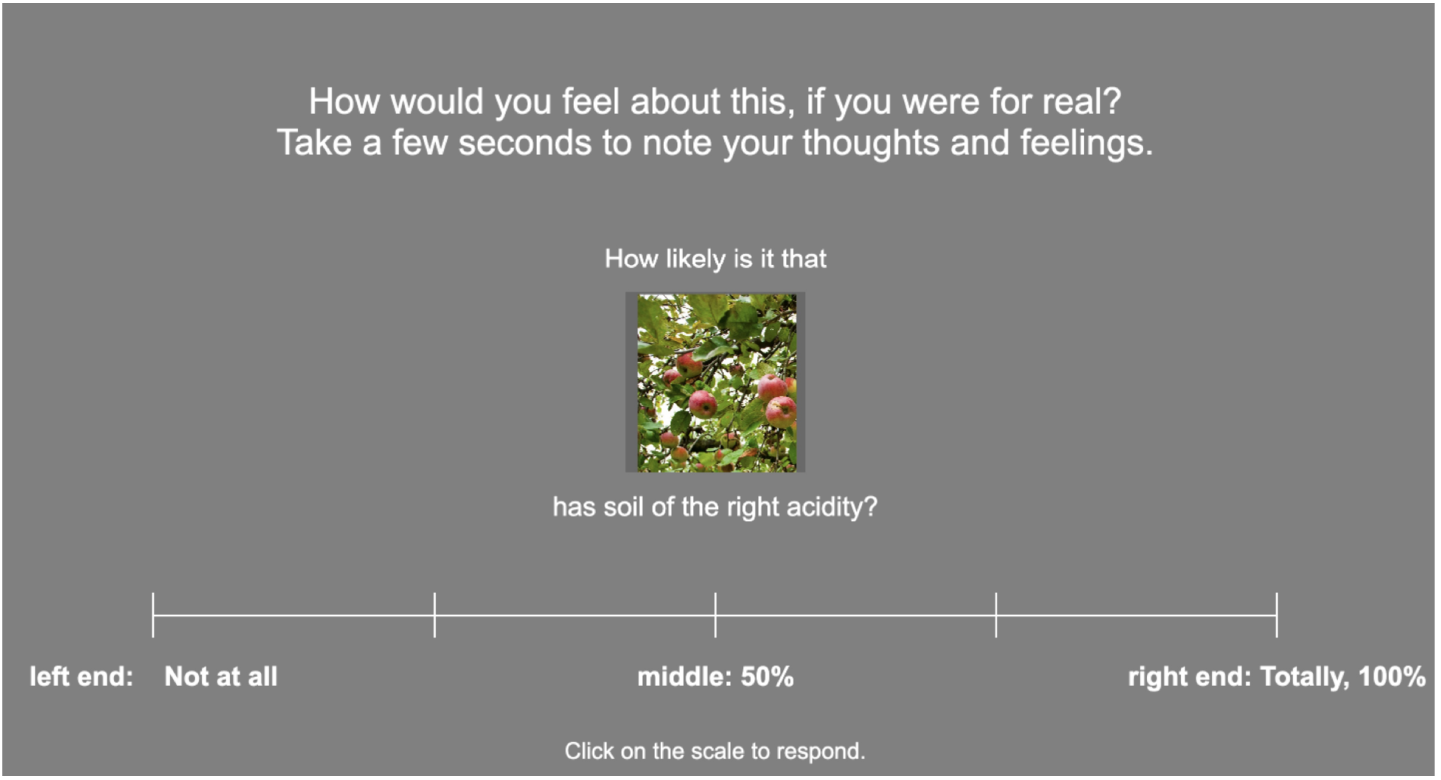
Sample screen-shot from the non-social, ‘apple trees’ task, at the point where the participant is asked for the attribution rating. This is the equivalent of being asked what they think the probability of the hotelier being caring is in the social task. Note that, like in the social version of the task, questions were always made in a positive frame – about the probability of the soil being good, the hotelier being caring, the apple tasting great, etc. This increased comprehensibility, but may have biased participants towards thinking primarily in terms of rewards and caring, rather than losses and uncaring hoteliers.

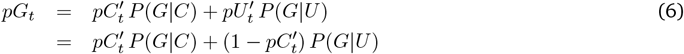

They then observe the outcome of trial *t* and update their beliefs about whether the underlying cause was beneficent, i.e. the hotel owner was caring. In the model, it is straightforward to use exact Bayesian updating:

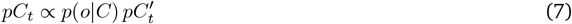

Participants do not report their beliefs veridically. They apply a bias *B*, which exponentiates the reported probability towards 1 if *B >* 1 or towards 0, if *B <* 1; Then, they respond noisily, subject to an inverse-temperature-like parameter *β* and a lapse rate, or noise floor, *ζ*. In the real experiment participants used visual-analogue scales, but for the purposes of modeling, we binned these into *n*_*r*_ = 6 equal response bins. It then becomes very convenient to use the escorted binomial distribution to model responses (Bercher, 2011). Let us denote the escorted binomial probability for *j* out of *n* successes, with success probability *q*, and order (here, inverse temperature) *β* as *EB*(*j*; *q, n, β*). We will use this to model a response in VAS bin *j*. We then derive the equation about the caring attribution response policy, *πC*_*t*_(*j*), in two steps, as follows (the outcome expectation response equations have exactly analogous form):

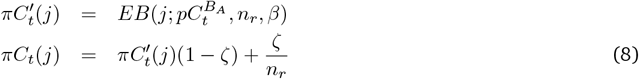

#### Valence-Partitioned Associative model (VPAL)

The value-update scheme for this followed very closely the valence-partitioned reinforcement learning formulation of Sands et al. (2023), as per eq 2. We described each trial as four states, with appetitive and aversive values learned for each. In the present, more formal description, it is clearer to use ‘+’ (rather than *P* in the main figures and text) to denote the Positive, or appetitive, channel, and ‘*−*’ ( rather than *N*) for the aversive channel. Gains *r*^+^ and losses *r*^*−*^ only entered the update of the appetitive and aversive channel respectively, and are each *≥* 0 by convention. Otherwise, they were treated as no-outcome. For example, if only a loss *r*^*−*^ were observed upon transitioning from state *s*(*k*) → *s*(*k* + 1):

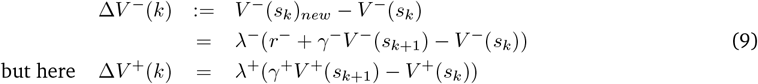

Our model differs from Sands et al. (2023) in two aspects only. First, it considers as terminal only the end state of each whole block. Otherwise, the transition *s*(4) **→** *s*(1) updates *s*(4). Secondly, the bias parameters above were re-purposed, to translate the reward and punishment TD values into belief-like fractions of the VAS scale offered. For example, in the case of Caring attribution self-report, we write for the fraction *f C* :

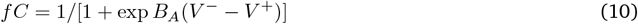

Crucially, this means that people’s psychological beliefs are not updated based on an explicit generative model. Here, it is a mathematically phenomenological input-output function, mapping reward expectations to psychological phenomenology. Also note that the ‘bias’ parameters, *B*_*A*_ etc. no longer bias in the sense of favouring one end of the scale rather than other, but function like extreme-responding or sensitivity biases, amplifying or blunt the difference between associative values in the map to phenomenological beliefs.

Finally, these belief-like fractions of the VAS scale are translated to actions using a policy *ϕ* very similar to *π* above :

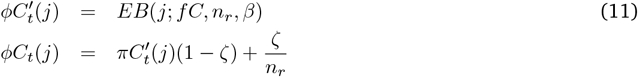

#### Temporal-Difference Valence-Partitioned Bayesian model (TD-Bayes)

The TD-Bayes model propagates beliefs through its four states exactly like the VPAL model propagates reward values; the difference here is that *V* ^+^ and *V* ^*−*^ do not represent reward and loss expectations, but log unnormalized beliefs about the beneficent or maleficent underlying cause.

The update of the prior probability for each partner proceeds as per eq. 3, 4 as above. However, now we apply the forgetting or volatility to the beneficent and maleficent cause channels, instead of eq. 5. For trial *t* of block *k*:

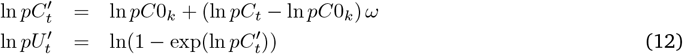

We now apply eq. 1 for the values *R*^+^, *R−* representing the un-normalized log probabilities at the point of observing the outcome:

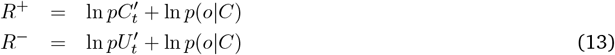

An interesting viewpoint, not explored here, would be to consider these two equations as evidence accumulation. One might then consider a further, attentional parameter weighting the log-likelihood term, which might desrcribe participants that do not pay attention to the observations (or even over-weigh them inappropriately).

## Additional modelling results

As our starting hypothesis was that valence-partitioned associative models might be used by the brain as useful approximations to Bayesian reasoning, we wanted to know whether the latent variables of these models, which had conceptual parallels, were distinguishable in practice. We examined, first, the decision-variables that formed the input into the response module of each model. These should correlate highly between VPAL and TD-Bayes, as behaviour was explained well by both models, and only response mechanisms with monotonic biases mediate between these overall-value or probability decision variables and the responses. Second, we examined the correlation of values carried by each of the positive or negative channels across models, noting that different combinations of such values may give rise to the same magnitude of the decision-variable (and hence have different brain signatures). Third, we examined correlations between the value- or belief-updates (the weighte prediction errors) across models.

**Figure S2:**
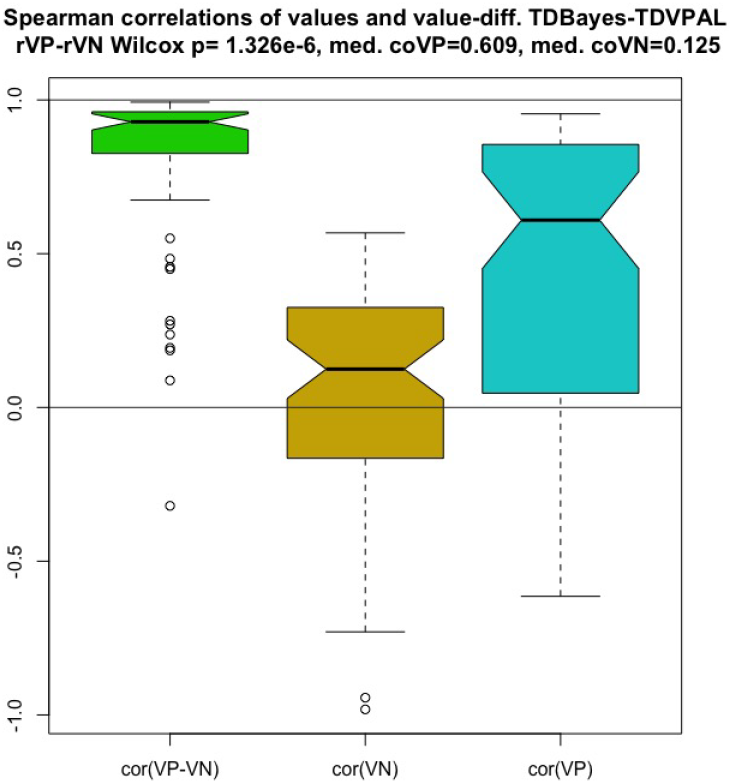
Relationship between latent measures of the two models for the two values, aversive and appetitive, and for the decision-variable, here the difference between the other two. A Spearman correlation was calculated for each participant and measure, and here the distribution across participants is shown. Notches are the standard errors of the median. Decision-variable measures are very highly correlated, as expected, but *V* ^+^/VP moderately so, and *V* ^*−*^/VN weakly so.

We found that, as expected, there was a very strong correlation between the decision variables, ie. the difference between the values of the appetive and aversive channels. The median-over-participants Spearman correlation for *V* ^+^ *− V* ^*−*^ being 0.928 (Fig. S2, *cor*(*V P− V N*)). The median correlation for *V* ^+^ was *cor*(*V P*) = 0.606 while for *V* ^*−*^ was only *cor*(*V N*) = 0.124, significantly positive but also significantly less than *cor*(*V P*). Their ranges had large overlaps (Fig. S2, *cor*(*V P*) and *cor*(*V N*)). These results mean that the models do produce different time-series of their latent variables, that neuroimaging can interrogate.

The correlations between updates showed a surprising pattern, with the aversive updates being significantly anti-correlated, albeit modestly so, median *cor*(*DV P*) = *−* 0.365. This may be because middle, ‘So-so’ outcomes constitute unambiguous (though quite modest) evidence for the aversive channel, according to the likelihood map that participants were trained on, but their associative reward value is not unambiguous – the ‘so-so’ outcome was taken to have neutral reward value.

**Figure S3:**
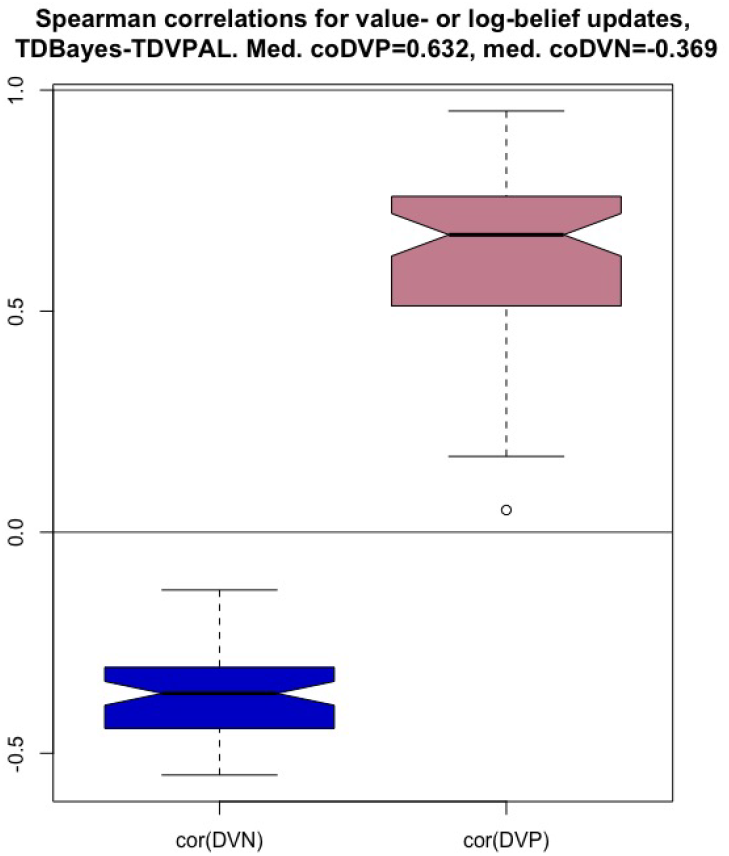
Relationship between latent belief- or value-update measures for the two models. The appetitive-update measures are still positively correlated, but the aversive-update measures negatively so.

